# Vertex-wise characterization of Non-Human Primate cortical development with prenatal insights

**DOI:** 10.1101/2021.09.23.461551

**Authors:** Julian S.B. Ramirez, Robert Hermosillo, Elina Thomas, Jennifer Y. Zhu, Darrick Sturgeon, Emma Schifsky, Anthony Galassi, Jacqueline R. Thompson, Jennifer L. Bagley, Michael P. Milham, Oscar Miranda-Dominguez, Samantha Papadakis, Muhammed Bah, AJ Mitchell, Ting Xu, Alice M. Graham, Eric Feczko, Elinor L. Sullivan, Damien A. Fair

## Abstract

Characterization of the interwoven complexities of early cortical thickness development has been an ongoing undertaking in neuroscience research. Longitudinal studies of Non-Human Primates (NHP) offer unique advantages to categorizing the diverse patterns of cortical growth trajectories. Here, we used latent growth models to characterize the trajectories of typical cortical thickness development in Japanese macaques at each cortical surface vertex (i.e. grayordinate). Cortical thickness from 4 to 36 months showed regional specific linear and non-linear trajectories and distinct maturation timing across the cortex. Intriguingly, we revealed a “accumulation/ablation phenomenon” of cortical maturation where the most profound development changes in cortical thickness occur in the accumulation or ablation zones surrounding the focal points (i.e., a center of a delineated regions where cortical thickness is thickest or thinnest) throughout the brain. We further examined maternal diet and inflammation in the context of these typical brain trajectories and known network architecture. A well-controlled NHP model of a maternal “Western-style” diet was used alongside measures of inflammatory cytokine interleukin-6 (IL-6) in the mothers during gestation. We observed that these accumulation and ablation zones of variable change might be most susceptible to environmental effects. The maternal factors, diet and inflammation during pregnancy were distinctively associated with different aspects of offspring cortical development reflected in regions related to distinctive functional networks. Our findings characterize the versatile intricacies of typical cortical thickness development and highlight how the maternal environment plays a role in offspring cortical development.

## Introduction

Cortical thickness measured with MRI has long been used to characterize the nature of the developing brain ^1–4^. Cortical thickness has been linked to several neurodevelopmental disorders, including attention-deficit / hyperactivity disorder (ADHD) ^5–10^, autism spectrum disorder (ASD) ^11–17^, schizophrenia ^18–20^, and in Alzheimer’s disease ^21^. Furthermore, the impact of genetic and environmental influences are often studied in the light of cortical thickness ^11, 22–27^.

### 1.1 Non-Human Primate models can supplement human studies of cortical development

While the underlying neurobiology supporting cortical thickness remains under investigation ^28, 29^, non-invasive MRI neuroimaging methods have made it possible to measure cortical thickness by calculating the width of cortical gray matter in surface space. This gray matter is made up of the neuronal bodies, and cortical thickness has been postulated to be related to the synaptic density, myelination and synaptic pruning of the cortex ^30^.

Though the utility of studying typical and atypical development of cortical thickness is well supported using MRI, accurate characterizations of human (particularly early) developmental models remain a challenge for several reasons. First, it is recognized that within-subject, longitudinal designs are critical for a proper definition of long-term growth trajectories ^31^; however, due to the protracted nature of human brain development, few longitudinal studies exist that capture the critical developmental window from birth through adolescence. In addition, it is well documented that increased subject motion in youth complicates longitudinal data collection and analysis through subject/data loss ^32^ and systematic image artifacts and acquisition complications ^33–39^. Other ‘interacting’ factors common to all human studies related to demographics, education, diet, and other ‘covariates’ also complicate the design of well-controlled longitudinal studies that measure cortical thickness.

Non-human primate models provide an opportunity to overcome some of these limitations of human longitudinal studies of cortical thickness. The developmental period from birth to early adolescence is truncated relative to humans allowing for repeated measurements to be acquired more easily. NHPs often undergo MRI while sedated and, thus, motion artifacts are limited relative to their human counterparts. In addition, environmental factors and timing of the data acquisition can be controlled. NHPs, to a far greater degree than rodents, share many complex behavioral and brain characteristics with humans ^40–49^. Last, NHPs have a similar gestational period to humans, with development of foundational brain organization predominantly occurring prenatally ^50^.

A few studies have already demonstrated the ability to capture components of cortical brain development from birth through early adolescence in macaques ^51–54^ and marmosets ^55–57^. However, they primarily focus on average cortical volumes instead of thickness in surface space and require a specific parcellation schema of the cortex. While these studies provide an important context for this report, our findings suggest key developmental characteristics may be missed when not investigating cortical thickness development in surface space on a vertex-wise scale.

### 1.2 Maternal factors may influence typical brain development

While macaque models provide an avenue to study longitudinal trajectories in typical development, they also offer several benefits when studying variability in development. Along with the factors noted above, the ability to implement well-controlled experimental designs with regard to various environmental factors has made NHP models tremendously advantageous ^50, 58–61^.

We have shown in several studies that maternal diet and inflammation during pregnancy can influence the offspring’s brain after birth. Human studies have found that maternal plasma interleukin-6 (IL-6) concentration relates to differences in growth characteristics of amygdala volumes and amygdala structural and functional connectivity ^62–65^. In macaques, maternal IL-6 is correlated with decreased amygdala volumes at 4-months, but also an increased rate of growth from 4 to 36-months-of-age, with an indirect relationship to anxiety-like behavior through alterations in amygdala development ^58^.

Other findings have demonstrated that a maternal “Western-Style” diet, high in fat, is associated with dysregulated dopamine and serotonergic systems in NHP offspring ^59, 66^.

### 1.3 Typical and atypical cortical thickness development in a macaque model

In the current report, we sought to characterize the typical developmental trajectories in cortical thickness from birth to early adolescence (36 months of age) in a Japanese Macaque model. To accomplish this, we investigated cortical change over four time-points in offspring development (Figure 1A). We utilized Latent Growth Curve (LGC) analyses ^67–70^ to identify the best-fitting models that characterize different growth trajectories on a grayordinate scale (Figure 1B). After establishing the unconditional developmental models, we also examined how these trajectories might be modified by maternal diet and inflammation (i.e., IL-6) while controlling for offspring sex. For this, we introduced the predictors and covariates (maternal IL-6, maternal diet, offspring sex) to the best-fitting latent growth variables for each grayordinate (see Methods section for complete analyses description). These methods enabled us to characterize typical macaque offspring cortical thickness trajectories and investigate how these may be impacted by maternal diet and maternal inflammation during gestation.

**Figure 1:**
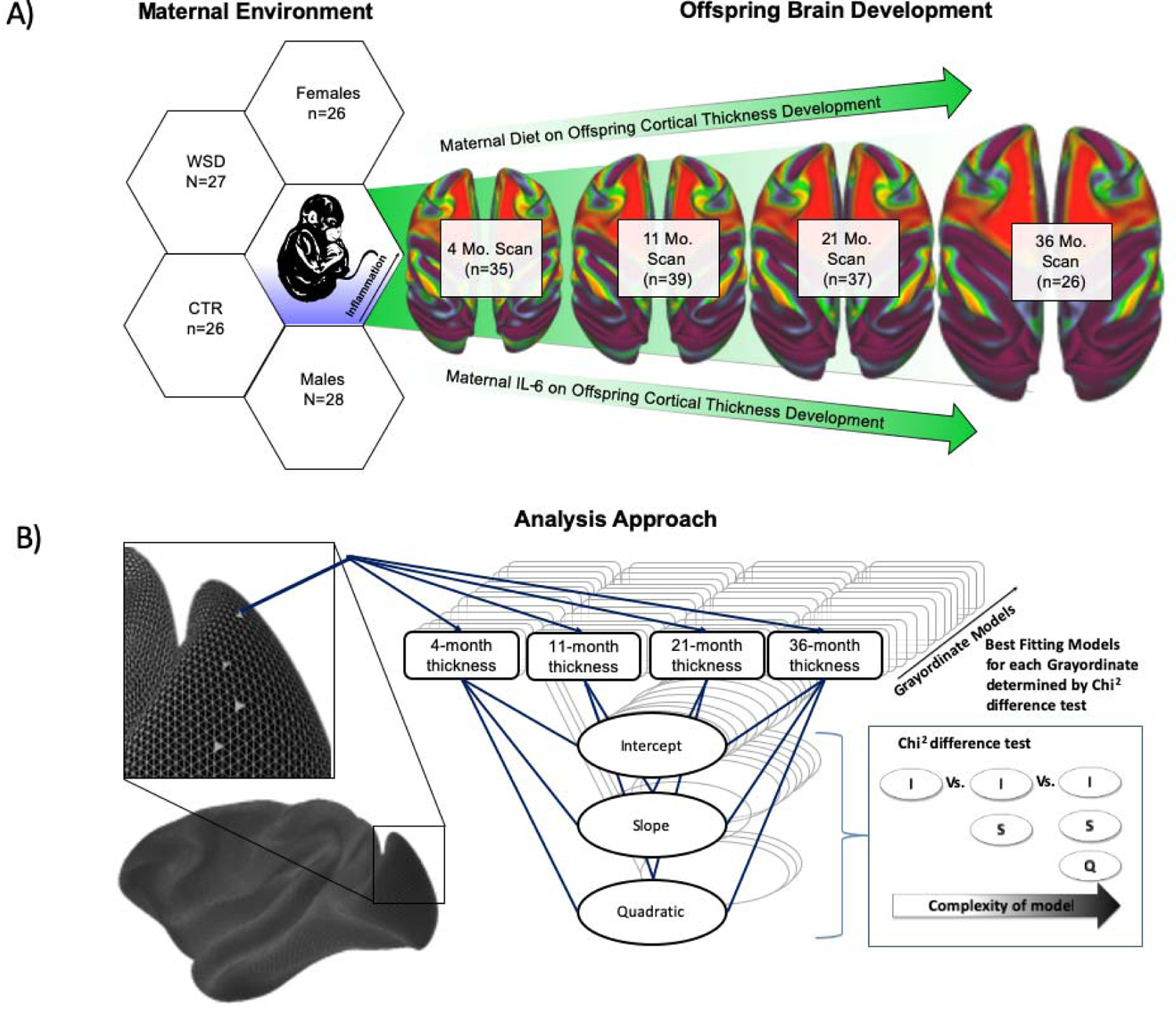
**A**: Study overview. Maternal Western-Style Diet (WSD) or Control Diet (CTR) and maternal IL-6 may influence cortical thickness development across time. **B**: Analysis model overview. Latent growth curve models are defined for each grayordinate, indicating the cortical thickness at 4, 11, 21, and 36-months of age. The simplest model (intercepts only) is run first, and then a Chi-Squared difference test is run on more complex models to identify the best-fitting model for each grayordinate.

## Results

### 2.1 Characterizing typical cortical thickness development across the entire brain

Here we characterized typical cortical thickness development using **1)** the best-fitting growth models (Figure 2A), **2)** the age of the monkeys at which the cortical thickness was thinnest (Figure 2B), or **3)** thickest at a particular region (Figure 2C), **4)** the estimated mean cortical thickness (Figure 3A-D), and **5)** the percent change in cortical thickness across time (Figure 3E-H).

**Figure 2:**
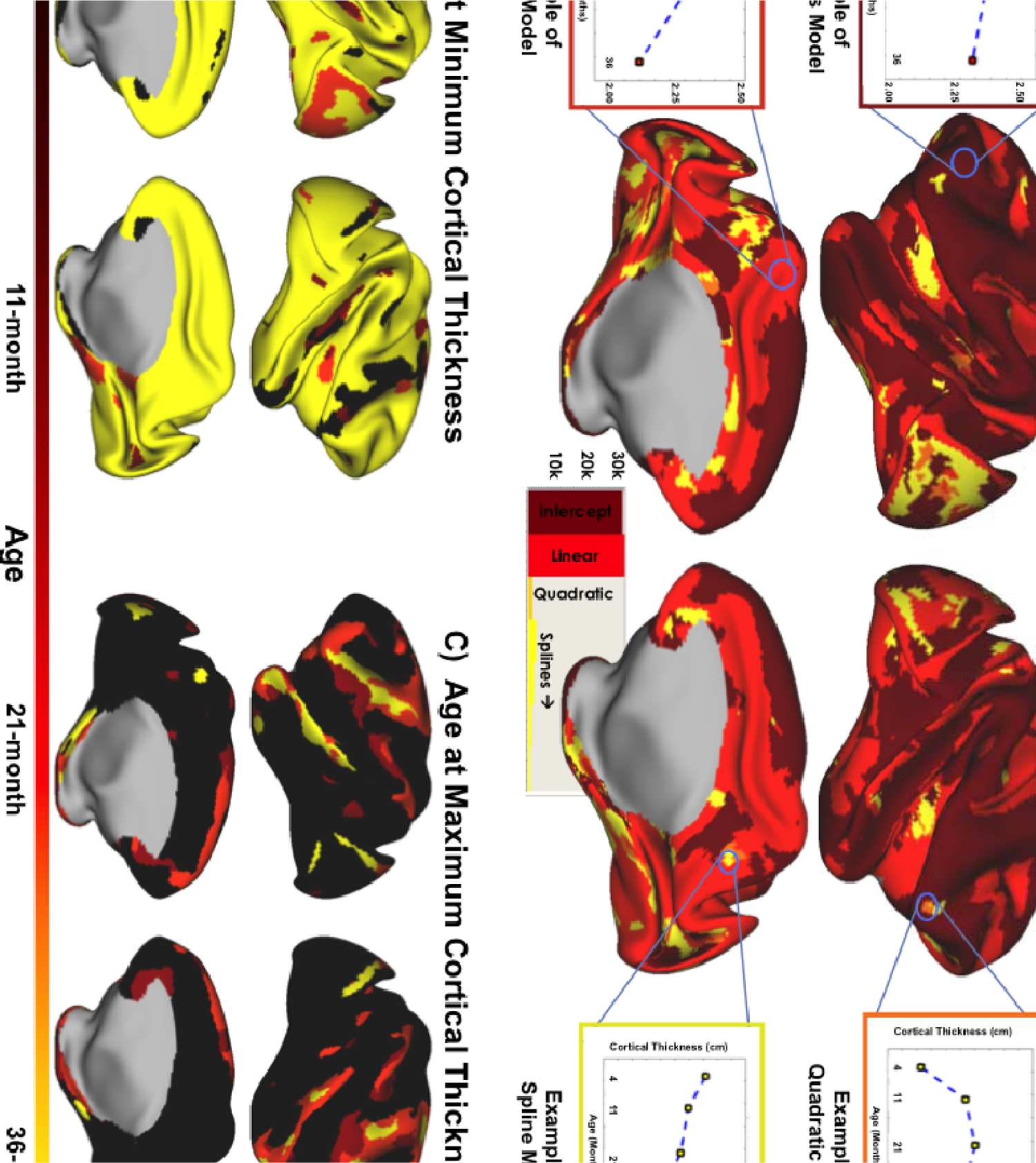
Best-fitting models are defined for each grayordinate using the chi-squared difference test comparing simple to more complex models. The best-fitting models are mapped onto the brain (A). A graph of mean cortical thickness for specific clusters where an intercepts-only (dark red), a linear (orange), a quadratic (light orange), or a spline (yellow) model best fit, to show the model fits represent actual development. (B) Shows the age at which the cortical thickness was thinnest. Regions where it is yellow, for example, indicate that cortical thickness was thinnest at 36-months of age. (C) Similarly, this shows the age at which cortical thickness was the thickest (max). Dark colors indicate that the maximum thickness was observed at 4-months of age. In cases where the thickness was the maximum at 4-months (dark red) and the minimum at 36-months (yellow), a gradual thinning over time can be inferred. Orange denotes an interesting pattern where a change in direction of thickness happened over time.

**Figure 3.**
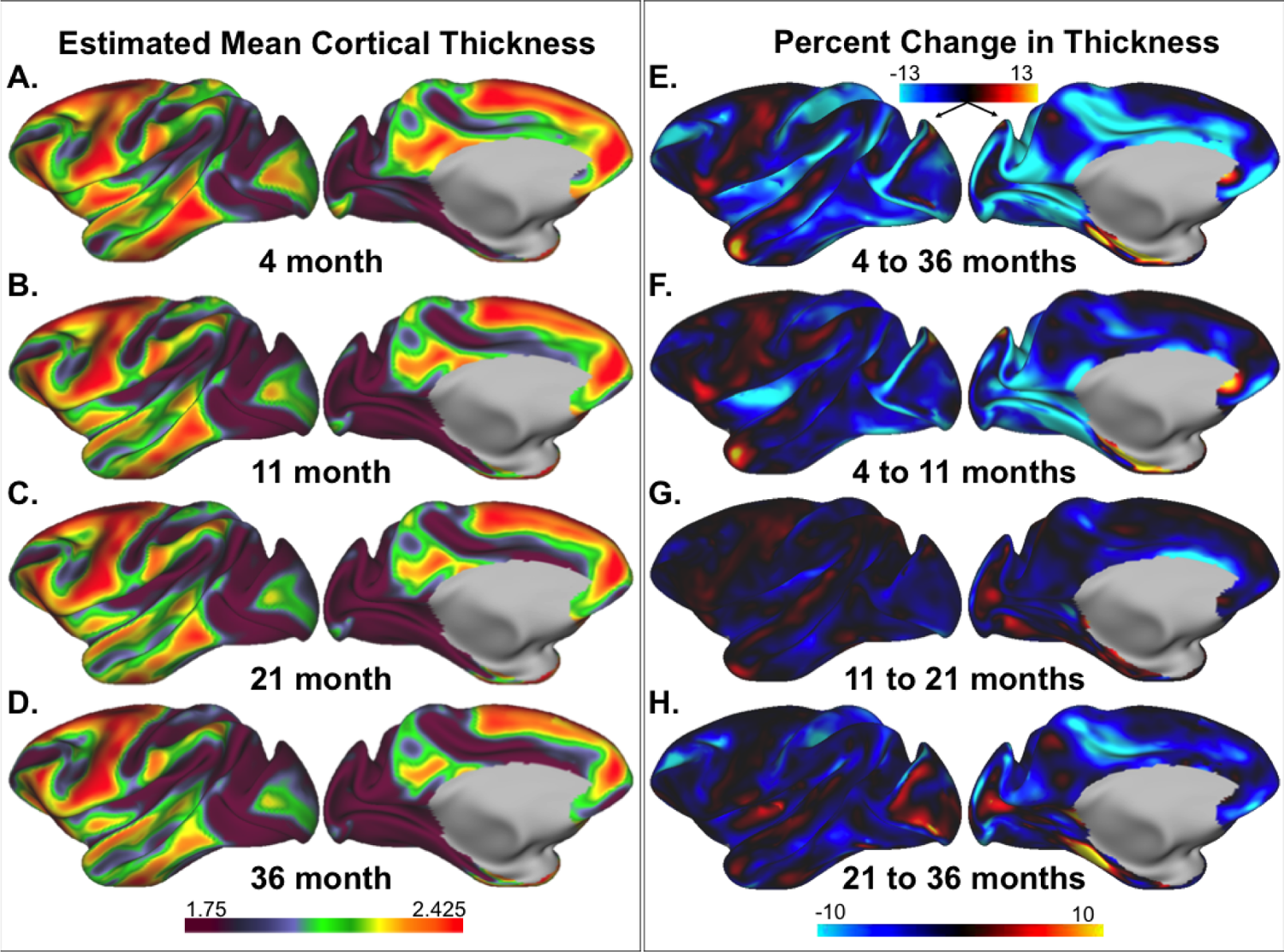
Cortical thickness development. The estimated mean thickness for different age groups (A-D) and percent change in thickness between 4 and 36-months (E) and more gradual stages in change from 4 to 11 months (F), 11 to 21-months (G), and 21 to 36-months (H).

#### The best-fitting growth models

We first aimed to characterize typical cortical thickness development for all cortical grayordinates using unconditional models. The best-fitting model for each grayordinate was identified by comparing intercepts-only, linear, quadratic, and spline models, choosing the simplest model that was not significantly improved via a more complex model (Figure 1A). The different growth patterns were mapped on the brain (Figure 2A). While the majority of regions in the brain were best characterized by either an intercepts-only model (i.e., little change across time) or a linear model (intercept and slope), some region’s development was best described by more complex growth models, including quadratic or spline functional forms (Figure 2A). This map of best-fitting models can be viewed alongside the supplemental movie 1.

#### Age at thinnest and thickest cortical thickness

We further characterized the age of the monkeys, when the cortical thickness was thinnest (Figure 2B) and thickest (Figure 2C). Not all regions start as being the thickest at the 4-month time point or end as being the thinnest at the 36-month time point. Instead, in some areas (e.g., the retrosplenial cortex, insular region, or the fronto-orbital gyrus), cortical thickness is thinnest/thickest at the 11 or 21-month time point – consistent with the non-linear best-model fits.

#### Mean cortical thickness

Using the unconditional model, we estimated and visualized the mean cortical thickness of each grayordinate across the four time-points (4, 11, 21, 36 months). The estimated mean cortical thickness decreased in the majority of the cortex across time (Figure 3). However, in some regions, the estimated mean cortical thickness increased from 4- to 36-months of age (Figure 3). Cortical thickness changed anywhere between −20 and 20 percent depending on the region (Figure 3E).

#### Percent change

When investigating the percent change between each time point, the most change occurred between the 4- and 11-month time points (Figure 3F), with the least amount of change occurring between 11- and 21 months of age (Figure 3G). The rate of cortical thickness development was not uniform across the brain. For example, while cortical thickness decreased ∼15% in insular regions between the 4- and 11-month time points, it increased from 21- to 36-months of age (Figure 3F & Figure 3H). This non-linear development can be seen more clearly in the supplemental movie 1 showing the estimated mean cortical thickness development (Supp. Mov. 1) and Supplemental Figure 2 showing both hemispheres (Supplemental Figure 2).

These above five different measures (i.e. trajectory models, age at thinnest, and thickest thickness, mean thickness, and percent change) utilized to characterize typical cortical development can help highlight unique and complementary components of cortical thickness development when considering a given region across these measures. For example, a pattern of non-linear change in the insula can be identified across approaches. The percent change in cortical thickness of the posterior insula (Figure 3F-H) shows an early decreased (Figure 3F) followed by a later increased (Figure 3G-H) percent change. This change in directionality of development was also captured by our best-fitting model approach in Figure 2A, showing a non-linear (yellow) best-fitting model in this region. The non-linear development of the insula was also reflected in Figure 2B. Here the minimum cortical thickness was reached around the 11 and 21-month time points (Figure 2B), indicating that cortical thickness must have plateaued or increased after 21-months of age. These are just some examples of how these different modalities can describe the changes that occur in cortical thickness as the brain develops. These measures could lend as a framework to guide future studies investigating developmental trajectories.

### 2.2 Vertex-wise analyses reveal dynamic ablation and accumulation zones in cortical development

It is important to note that different areas in the brain do not uniformly change across time. More interestingly, this change occurs in gradients in zones surrounding the focal points of peak thickness or thinness regions (i.e. accumulation/ablation zones, Fig 4A), instead of the center of the peak regions. These findings highlight the use of fine-grained developmental models at vertex-wise (i.e., grayordinate) scale, which enables characterization of the gradient patterns in growth trajectories along the cortex that would be missed using predefined regions of interest (ROIs) or an areal parcellation scheme.

**Figure 4:**
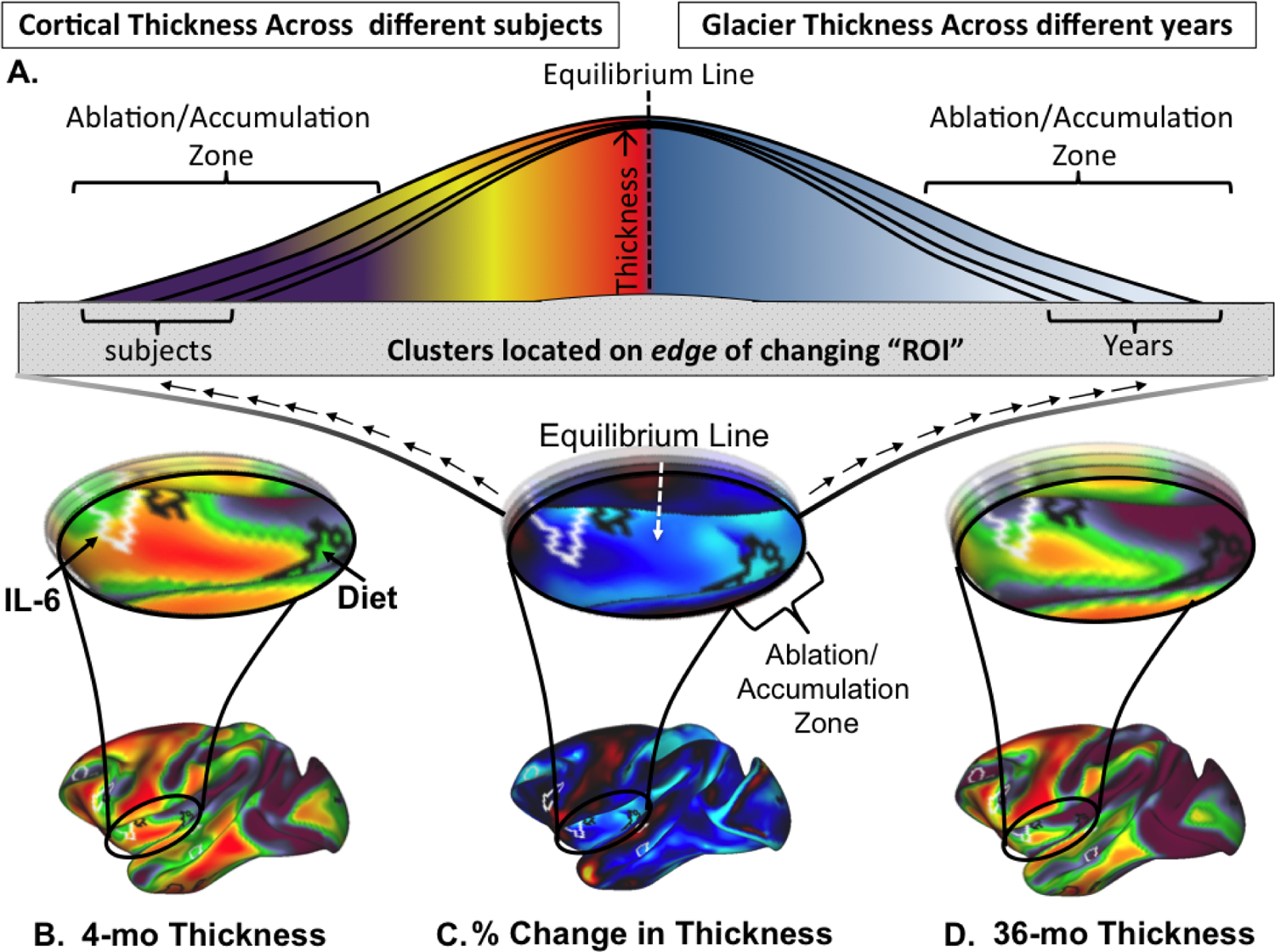
Impressionistically, many clusters fall on the edge of focal points where max or min thickness occurs, rather than at the center. We describe this phenomenon by analogy to glacier melt. Rather than seeing the most change in melting in the center, this occurs on the edges (i.e., the ablation zone). These phenomena are similar in the current findings and glacial research.

Here we describe the zones of gradient change surrounding a focal point by analogy to glacier melt in nature. The glacial melt across time most prominently occurs on the edge (i.e., the ablation and accumulation zones) as opposed to the center of the glacier (i.e., focal point or equilibrium line) ^71^. We observed a similar phenomenon in our findings of cortical development, highlighting multiple focal points with surrounding “ablation” and “accumulation” zones throughout the brain. Borrowing from the glacial literature, this area of gradual change around a focal point will be framed as the “ablation zone” (cortical thinning gradient) or the “accumulation zone” (cortical thickening gradient) for the remainder of the manuscript ^71^. This comparison is depicted in Figure 4A.

For example, in the insular region, the cortical thickness is at its thickest in the center of this ROI at 4-months of age (Figure 3A). Consider this point the focal point. As the monkeys get older, the cortical thickness decreases or ablates in the insula, however not in a uniform fashion. The thickness surrounding this focal point ablates at a different rate in a gradient zone around the center (Figure 3A-D). This phenomenon is also seen in Figure 2B, where the minimum cortical thickness is reached at 21-months of age in the center of the insula but at 36-months in the surrounding zone as cortical thickness continues to ablate. These zones of gradient change towards a focal point would be harder to observe using ROIs, as the whole insula ROI would be considered to get thinner uniformly over time.

### 2.3 Maternal diet influences cortical thickness development around accumulation/ablation zones

Having established the unconditional models describing typical cortical thickness development, next, as a proof of concept, the predictors (Maternal Diet, Maternal IL-6) of interest were introduced to identify how these related to unconditional developmental trajectories while controlling for offspring sex. Interaction terms were not added to maximize the convergence of the models. Maternal diet and IL-6 both had independent and overlapping relationships to the different latent growth variables (i.e., intercept, slope, quadratic) that characterized typical development. How the predictors are associated with increased or decreased growth variables can be examined on the brain showing the overall estimates (beta weights) or the most significant clusters for each latent variable (Figure 5 & Figure 6), as well as an example of where they fall in a given ablation zone (Figure 4B-D). Most significant clusters were identified through a cluster detection algorithm based on previously defined guidelines ^72^ which is described in more detail in the methods section.

**Figure 5:**
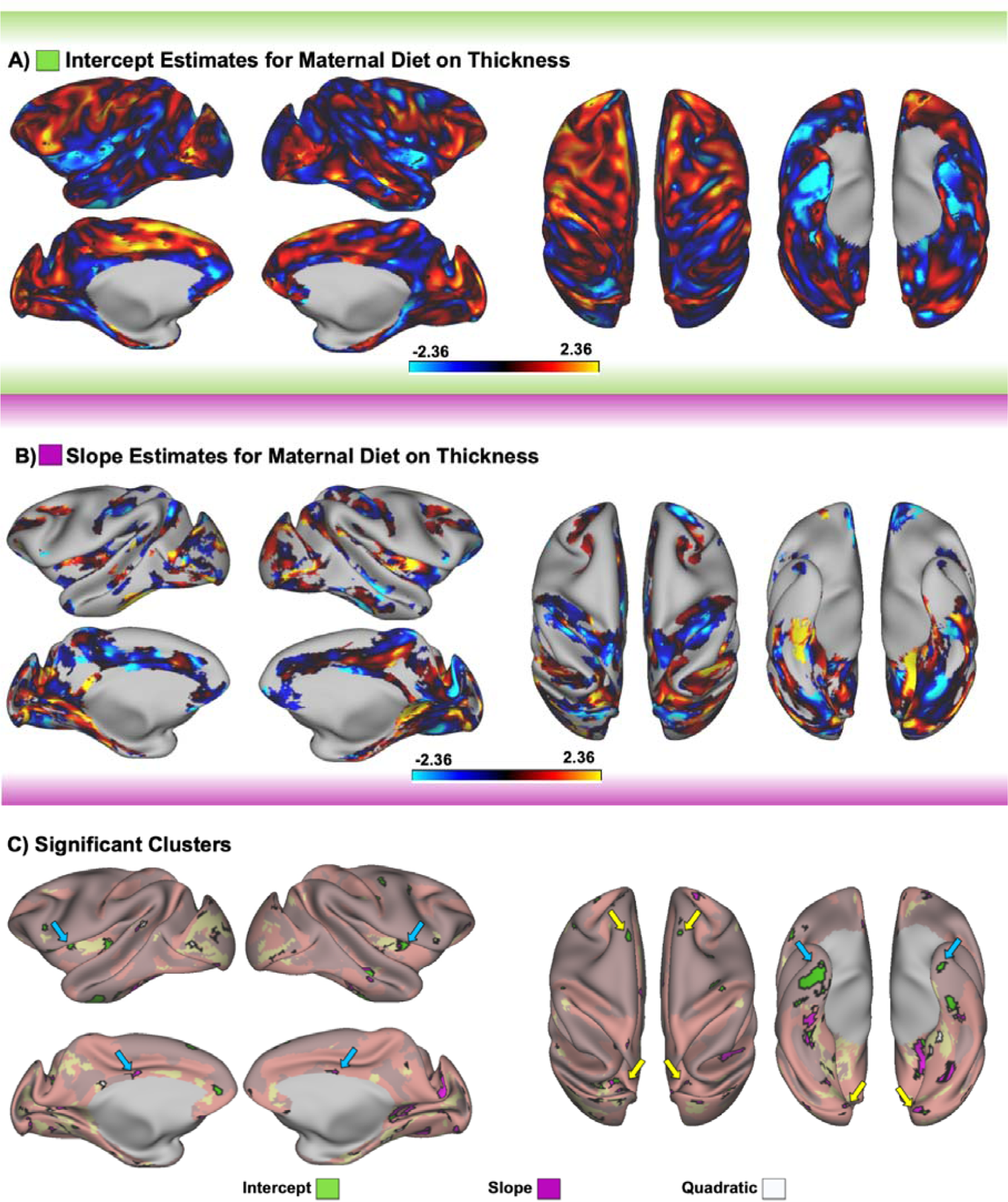
The influence of diet on cortical thickness development. (A) indicates the estimates of how diet relates to the intercepts of the models (B) shows the estimates of how diet relates to the slope parameter (in regions where an intercept only model was not the best fitting model), and (C) highlights the significant clusters from this analysis which are overlaid on a transparent best-fitting model map from figure 2. Green clusters are clusters where diet had a significant relationship with the intercept, and pink clusters show the significant slope clusters. White clusters indicate the quadratic; however, since there were so few best-fitting quadratic models, the estimates are reported in the supplemental materials. Arrows highlight some interesting bilateral clusters where a WSD was associated with decreased (blue arrow) or increased (yellow arrow) intercept (green cluster) or slope (pink cluster). The direction of association (highlighted by the color of arrows) can be identified by the color of the estimates (A&B).

**Figure 6:**
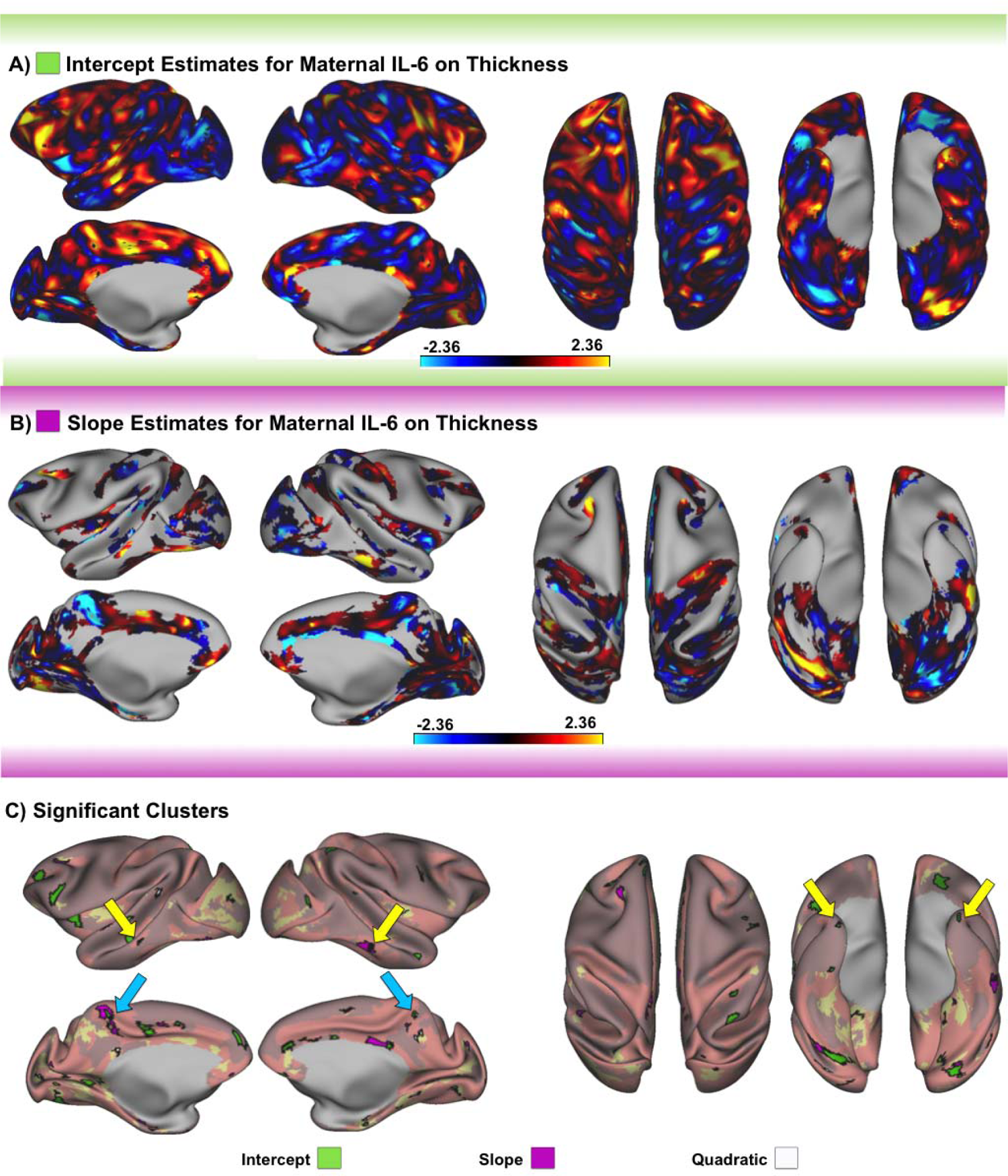
The influence of maternal IL-6 on cortical thickness development. (A) Indicates the estimates of how IL-6 relates to the intercepts of the models (B) shows the estimates of how IL-6 relates to the slope parameter, and (C) highlights the significant clusters from this analysis which are overlaid on a transparent best-fitting model map from figure 2. Green clusters are clusters where IL-6 had a significant relationship with the intercept, and pink clusters show the significant slope clusters. White clusters indicate the quadratic; however, since there were so few best-fitting quadratic models, the estimates are reported in the supplemental materials. Arrows highlight some interesting bilateral clusters where higher concentrations of IL-6 were associated with decreased (blue arrow) or increased (yellow arrow) intercept (green cluster) or slope (pink cluster). The direction of association (highlighted by the color of arrows) can be identified by the color of the estimates (A&B).

#### Intercept

Maternal Western-style diet (WSD) was associated with both increased and decreased cortical thickness at the intercept (i.e. 4-months) of several regions (Figure 5A & Figure 5C [green clusters]). Bilateral clusters highlighting where the WSD was associated with decreased (blue arrows; green clusters) or increased (yellow arrows; green clusters) cortical thickness early in life (intercept/4-month time point) emerged throughout the brain (Figure 5C).

A common theme in the results of the cluster analysis was that clusters tended to fall within the previously defined ablation and accumulation zones (Figure 4B-D & Figure 5C). This phenomenon can be clearly seen in the insular clusters of Figure 5C. These clusters fell in the ablation zone surrounding the insula’s focal point (Figure 4B-D). This phenomenon can also be depicted in four scenarios of Figure 2 & Figure 3, showing that the clusters would fall on 1) the edge of where the best-fitting model changed from a linear to a non-linear trajectory in the insula (yellow area in the insula of Figure 2A), 2) the edge of the local maxima of mean insular cortical thickness (Figure 3A), 3) the edge between the 21 and 36-month age points defining the minimum cortical thickness (Figure 2B) and 4) the area of the highest percent change in the insular region (Figure 3E).

#### Slope

Several clusters also showed that maternal WSD was associated with an increased or decreased rate of growth over time (Figure 5B & Figure 5C [pink clusters]). Potentially interesting bilateral clusters showing the association between maternal WSD and a decreased (blue arrows; pink clusters) or increased (yellow arrows; pink clusters) rate of cortical thinning are highlighted in pink in Figure 5C. There were also instances where a slope cluster overlapped with an intercept cluster showing an opposite direction (increased or decreased) of association (e.g., the right posterior insula). This would indicate that WSD was associated with a decreased initial cortical thickness (intercept) and an increased rate of cortical thinning (slope) in this example. This type of bi-directional relationship is consistent with our recent findings relating maternal IL-6 to amygdala volume development ^58^.

#### Quadratic

As there were so few regions that were best characterized by a quadratic, for completeness, these data are included primarily in the supplement (Figure 3A & Supplemental Figure 3). No notable bilateral clusters were identified for this measure (Figure 5C [white clusters]).

### 2.3 Maternal IL-6 showed unique and overlapping relationships to cortical thickness development as compared to maternal diet

#### Intercept

As with maternal diet, IL-6 also showed increased and decreased relationships with the intercept (Figure 6A & Figure 6C [green clusters]) depending on the brain region. While IL-6 and maternal WSD showed similar effects in some regions, in other regions opposing effects were observed. For example, higher maternal IL-6 was associated with increased bilateral 4-month cortical thickness in the pole of the temporal cortex (Figure 6C [green clusters; yellow arrows]), which is an opposite effect of what was observed from the maternal WSD temporal pole results (Figure 5C [green clusters; blue arrows]).

#### Slope

When IL-6 concentration was used to predict the rate of change in thickness (i.e. slope) (Figure 5B), we observed a significantly decreased (blue arrows; pink clusters) or increased (yellow arrows; pink clusters) bilateral clusters. These clusters are highlighted in Figure 6C (pink clusters). Like the WSD findings, no meaningful bilateral clusters stood out with the quadratic term (Figure 6C [white clusters] & Supplemental Figure 3).

Both maternal IL-6 and diet were correlated with region-specific changes in cortical thickness. Modeling mean whole-brain cortical thickness development with maternal diet and IL-6 indicated that mean-cortical thickness was too broad to capture the complex relationships between maternal diet, IL-6, and cortical thickness development (Supplemental materials & Supplemental Figure 4).

### 2.5 Contextualizing cluster findings in the context of networks highlighted the broad vs. confined impact of diet and IL-6, respectively

The resting-state fMRI field has demonstrated the importance of thinking about changes in the brain on a network level instead of a single region. Prior work has highlighted that NHPs have similar network organizations to humans ^43, 73–75^.

Contextualizing the cluster findings in the context of large-scale functional networks may add to the interpretation of these findings for translational purposes. This is of particular importance in the light of recent work focused on reproducible brain-wide association studies (BWAS) in humans ^76^. While the longitudinal nature of the current cohort allows for repeated measures which increases power with regard to this point, specific cluster locations may not be directly generalizable due to the relatively small sample size of the current cohort.

While each individual cluster may not be drastically informative, the sum of multiple clusters may highlight important regions or functional networks that may be altered by maternal diet or IL-6. Consequently, after establishing the significant clusters associated with the predictors and cortical thickness development, the surface area of these clusters was calculated and compared using a chi-square test based on which functional network they fell into ^74, 77^.

Maternal diet was related to differences in cortical surface across a broad number of functional networks. The most robust differences associated with maternal diet were observed in the Default Mode Network (DMN), Visual (VIS), Auditory (AUD) and the Somatomotor (SMN) networks. The finding that the default mode and visual networks show similar trends is consistent with prior work by Power et al, highlighting that these systems tend to ‘hang’ together with regard to network characteristics ^78^. Specifically, there was a higher than expected cluster surface area in the DMN (χ^2^ (1) = 25.29, p >.001) and a lower than expected surface area in the Limbic (LIM) (χ^2^ (1) = 6.32, p >.05), Dorsal Attention (DAN) (χ^2^ (1) = 11.00, p >.001) and SMN (χ^2^ (1) = 7.92, p >.01) networks for the diet intercept (4-month) results. Furthermore, the surface area was higher than expected in the VIS (χ^2^ (1) = 46.90, p >.001) network and lower than expected in the LIM (χ^2^ (1) = 5.08, p >.05), DAN (χ^2^ (1) = 9.64, p >.01), Insular Opercular (INO) (χ^2^ (1) = 6.33, p >.05) and SMN (χ^2^ (1) = 15.21, p >.001) for the diet slope (growth) results. Finally, quadratic results showed that diet had higher VIS (χ^2^ (1) = 20.44, p >.001) and AUD (χ^2^ (1) = 8.25, p >.01) with lower DMN (χ^2^ (1) = 4.03, p >.05) and SMN (χ^2^ (1) = 8.46, p >.01) surface area than expected (Figure 7).

**Figure 7:**
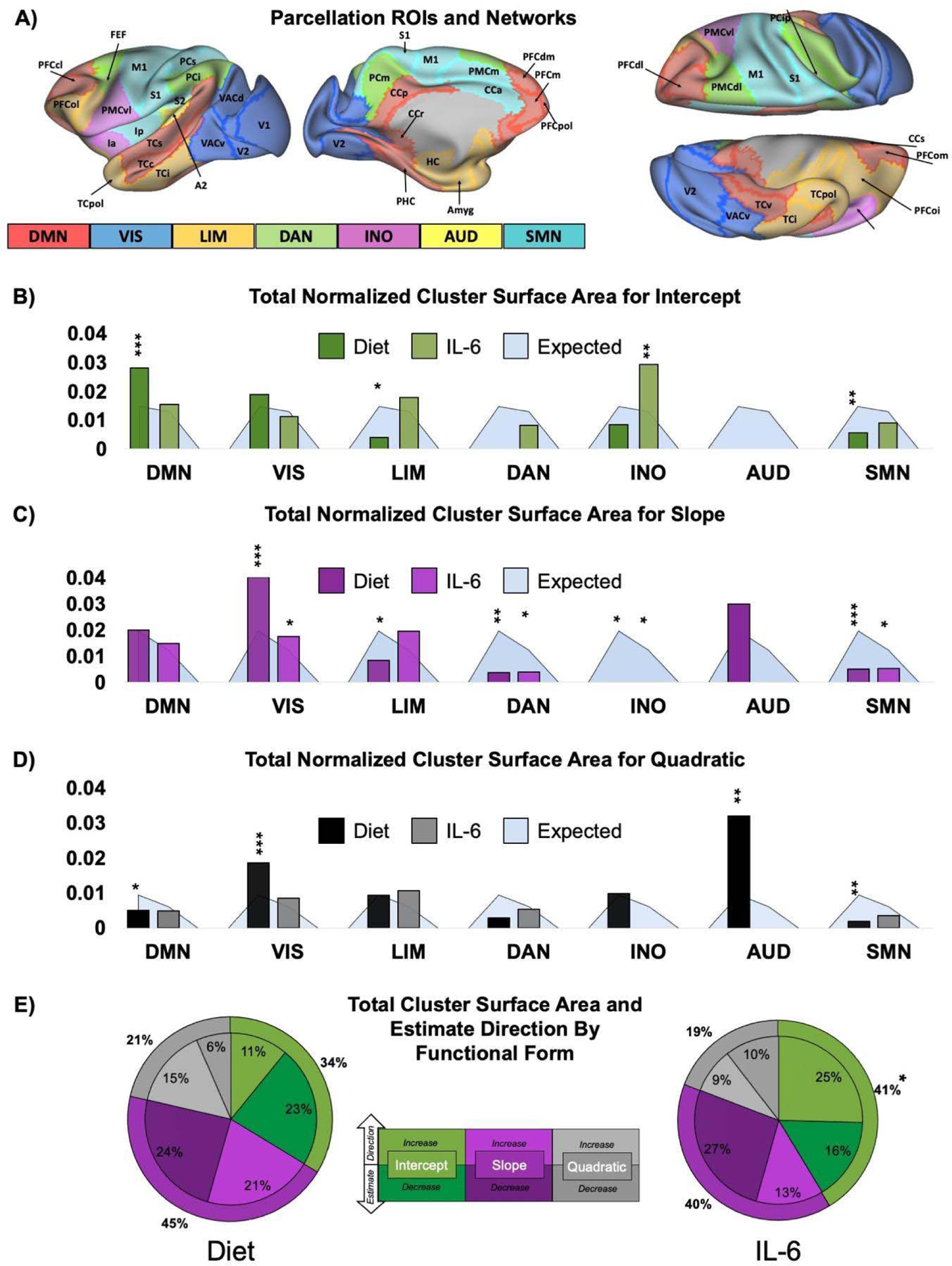
Summary results describing the cluster surface area trends. (A) Shows the functional networks and ROIs ^77, 79^. (B-D) highlight the surface area of the significant clusters for the intercept (B) slope (C) and quadratic (D). These graphs show the significant clusters for each predictor (Maternal Diet, IL-6) and where, in relation to the parcellation, these clusters fell, as indicated by the network they are in. The total cluster surface area is normalized by network size and plotted on top of the expected surface area for each network and predictor (light blue). Finally, the pie charts (E) indicate the percent of the surface area of the clusters related to the intercept (green), slope (pink), or quadratic (grey), with lighter colors indicating a positive (i.e., the predictor is associated with increased thickness) and darker a negative (i.e., the predictor is associated with decreased thickness) association. Note: P<0.05; P<0.01; P<0.001.

Maternal IL-6 was associated with a more confined network result, with the Insular Opercular (INO) network showing the most significant outcomes. Intercept IL-6 clusters were significantly higher in the INO (χ^2^ (1) = 6.80, p >.01) network than expected. Interestingly, slope results showed that maternal IL-6 was associated with significantly less surface area than expected in the INO (χ^2^ (1) = 3.98, p >.05) outcomes.

Similarly, IL-6 was associated with a reduction in network cortical thickness surface area of the DAN (χ^2^ (1) = 4.24, p >.05) and SMN (χ^2^ (1) = 5.76, p >.05), but increased surface area of the VIS (χ^2^ (1) = 4.84, p >.05) network (Figure 7).

## Discussion

### Overview of findings

The current report comprises a number of noteworthy insights into typical macaque cortical thickness development and how prenatal factors may relate to this characterized brain maturation. Specific growth models were needed to delineate developmental trajectories depending on the region of the brain. Cortical thickness between 4 and 36 months showed multiple trajectories, including very little change across time (intercepts-only model), consistent linear change across time (Slope models), or more complex non-linear changes across time (quadratic or spline models).

Examining typical trajectories on a grayordinate level was important as it revealed an interesting phenomenon that areas of dynamic accumulation or ablation in thickness development occur in zones surrounding multiple focal points throughout the brain.

We further examined the effects of maternal diet and inflammation on these typical brain trajectories and in the context of these precise dynamics and in the context of known network architecture. Novel observations indicated that many of the significant clusters associated with maternal factors fell within the accumulation and ablation zones, where typical changes over time were most dynamic. In addition to showing the susceptible nature of these zones, we also discovered that maternal associations manifested in a highly bi-directional fashion and were concentrated in meaningful patterns in relation to well-established macaque networks ^74, 77^. While diet showed widespread network associations throughout the brain with the strongest effects in the default, visual and somatomotor networks, maternal IL-6 showed a more focal relationship occurring predominantly in the insular-opercular network (Figure 7).

The strong bi-directional nature and network representation of our findings in a longitudinal model and our overall sample size of n=53 animals offsets some of the concerns that may arise regarding the reproducible nature of brain-wide association studies ^76^ using the smaller sample sizes that are unavoidable in non-human primate research. The decrease in life-style variability and controlled environment of a macaque model may also help offset some of the sample size issues. This study is unique since only a small number of longitudinal infant monkey MRI studies exist because these data are exceptionally rare and hard to attain, with only seven primate research centers funded by the NIH existing in the United States. For this reason, the sample size in this study is larger than most of longitudinal NHP studies of brain development. Further, this is the first study to examine the influence of a maternal “Western-Style” diet on brain development in NHPs.

### Typical macaque cortical thickness development highlights similarities and differences to the human literature

These results successfully characterized development and also mirrored broad growth trends observed in the human literature, lending to the translational value of using a macaque model. Visually inspecting the typical macaque change in cortical thickness over time highlighted a number of patterns similar to what is described in the human literature. For example, the increases observed in thickness of the motor cortex, the temporal pole, the parahippocampus, and some visual regions are also reflected in human literature, which shows either a comparatively lower rate of thinning or even increased thickening relative to other regions in the brain ^3,80–82^. Furthermore, regions considered to have the most change in cortical thinning over time in our NHP model, such as the medial and inferior parietal cortex, the centrolateral/pole of the prefrontal cortex, and the insula, all showed similar characteristics to the human literature ^3, 80–82^. The insula findings were less conclusive across these four studies, as only Amlien et al., 2016 showed this trend, with the other studies either not showing this data or having mixed trajectories. As the developmental time frame examined was different across these studies, this could relate to some of the disparities. For example, these studies ranged from 0-2 years ^82^or 4-10+ years of age, which is not directly comparable to the current study.

As few studies cover a similar timeframe as the current study (roughly one year through early puberty in human time), direct comparisons, especially with different trajectories, are difficult. However, a recent study that quantified typical cortical thickness development between the ages of one and six used a similar approach and found similar findings ^83^. In terms of percent change in thickness for the 68 (34 per hemisphere) regions looked at by Remer et al., the regions showing the most negative change (decreased thickness) also showed a similar direction and magnitude of change as we observed in the current study. Additionally, the regions with the most positive change (increased thickness) also overlapped with the patterns of change that we observed in the present findings. Furthermore, while this study only used linear, logarithmic, and cubic functional forms, there was some overlap in terms of our linear and non-linear findings as well. Naturally, the results were not directly comparable, as the current study used intercepts-only, slope, quadratic, and spline terms. However, in regions where Remer et al., used non-linear functions (quadratic or logarithmic), there was overlap in regions that were best described by our non-linear (quadratic and spline) functions ^83^. Generalized additive models (GAM) or Generalized additive models for location scale and shape (GAMLSS) could also be interesting models to test in longitudinal studies.

In order to have an even more accurate translational comparison across species, future work will require comparisons across similar developmental stages, using similar longitudinal statistics and precision grayordinate level analyses. Current work by ^75^, which allows for transformation between human and NHP models, would make such a study even more comparable. Ideally, these studies would characterize cortical development on a voxel, or grayordinate scale, where gradients of change are more easily observed. This is important given the findings of the current study which indicate that environmental factors like maternal IL-6 or diet may actually shift cortical trajectories in the ablation and accumulation zones surrounding the central points of a given ROI, which could be missed if only analyzing the trajectories of the entire ROI.

### Characterizing growth trajectories at the fine-grained grayordinate level showed that development might be most vulnerable in ablation and accumulation zones

Our typical results examining cortical thickness development indicated that developmental trajectories are not uniform across one region of interest. Modeling cortical thickness development on a grayordinate level proved to be beneficial, as it allowed us to identify the heterogeneous nature of brain development beyond the ROI level. The diverse nature of cortical thickness development can be characterized via multiple approaches such as describing the best-fitting model per grayordinate (Figure 2A), the age at maximum and minimum thickness (Figure 2B&C), and the mean and percent change in thickness across time (Figure 3). More importantly, running these analyses on a grayordinate level revealed ablation and accumulation zones, which modified our traditional view of how a variable may relate to differences in the brain.

When studying the impact a variable may have on the brain the initial assumption may be that this variable would most prominently affect the center of an ROI. The typical BWAS study will report findings in a way where one variable is linked to differences in a given ROI, without taking into consideration the developmental component of how this ROI is formed over time. We tend to look at changes in a particular region, such as the anterior insular (Figure 4B & D) and expect that the center, where mean cortical thickness is the greatest, is the area being affected by a given variable. However, the data here describe a different story by emphasizing the notion that group differences may be most prominently seen as a factor of the “edge” *development* of an ROI, rather than differences in the center of the ROI itself.

Indeed, when looking at the cluster results underlining the areas most likely to be impacted by maternal diet or maternal IL-6, we found that the center of a region may not actually be the place most vulnerable to change. Here, we observed that a number of prominent clusters fell in the zones surrounding these focal points rather than falling within the center of the region as one might expect. While this was unexpected at first, this observation fits into the glacier analogy framework explained above (Figure 4). In a glacier, between the accumulation (top of the glacier) and ablation (bottom) zones is the equilibrium line. As one moves further away from the equilibrium line towards the edge, much more rapid and variable accumulation or melting/evaporation of snow/ice is seen over the seasons and years making these zones more susceptible to environmental impacts ^71^. Analogously, this cluster phenomenon in the brain could be seen in the same light, as the brain is developing, more changes will be experienced on the edge (ablation zone) of the focal point (equilibrium line) of the ROI, explaining why the largest differences in cortical thickness by predictors were found in these regions. This is illustrated in (Figure 4). The current maternal diet and inflammation findings highlight the vulnerable nature of these accumulation and ablation zones and how differences in the maternal environment may alter the trajectory of how these ROIs are formed. Having a better understanding of the dynamic characteristics of cortical thickness development may help frame the foundation of existing literature describing how perinatal diet and inflammation may relate to brain function and structure.

### Maternal diet and inflammation cortical thickness findings supplement prior functional MRI studies

Although this is the first study to investigated the influence of a maternal Western-style diet or interleukin-6 on cortical thickness development specifically, research studying maternal factors such as inflammation or obesity have shown differences in structural and functional brain characteristics ^84–88^. On a network level, maternal IL-6 has been associated with within network connectivity of the subcortical, dorsal attention, salience, cerebellar, ventral attention, visual, cingulo-opercular, and frontoparietal networks, and between network connectivity of the subcortical-cerebellar, visual-dorsal attention and the salience-cingulo-opercular networks ^64^. Regarding the connectivity of specific ROIs, prior research has related inflammatory markers to increased connectivity of the left and right insula, the dorsal anterior cingulate, and the right and left amygdala ^63, 65^. Furthermore, these studies also showed decreased connectivity of the dorsal anterior cingulate as well as the left and right amygdala ^63, 65^.

Finally, inflammatory markers were also related to increased right amygdala and ventricular volumes as well as the rate of left amygdala volume growth ^58, 63, 89^. With an additional relationship indicating decreased entorhinal cortex, posterior cingulate, and left amygdala volumes ^58, 89^.

While these measures do not directly relate to cortical thickness, they do highlight some of the clusters found surrounding the insula, the cingulate, and some of the networks in which the clusters fell, such as our IL-6 insular-opercular network connection. Future research will need to investigate these relationships further; however, the current findings show regions that could be targeted for these types of studies.

#### Limitations

There are some limitations that need to be taken into consideration when interpreting these results. First, the sample size of this study, although larger than most NHP MRI studies, was much smaller than the typical sample sizes observed in human literature. Recent reports in humans highlight the importance of having an adequate sample size to identify reproducible brain-wide associations ^76^. For this reason, individual clusters should be interpreted with some caution and seen as potential areas to explore further. However, while the examination of the influence of maternal diet and IL-6 on the development of cortical thickness was performed as a proof of concept, it is important to note that the power of this study to detect an effect was enhanced by the repeated measures design. Furthermore, the number of regions that were consistent across the two hemispheres help validate the findings in a sense as a within-study “replication.” Additionally, in order supplement the BWAS limitation, and to highlight the broad patterns of interest rather than placing emphasis on the individual clusters, we combined all of the cluster results together, characterizing them into functional networks.

Despite these limitations these findings contribute valuable insights into typical development characteristics in a longitudinal within-subjects design. Nonetheless, running these types of growth models is still difficult with smaller sample sizes, as including multiple covariates in the model can cause the model not to converge or produce other statistical warnings. For this reason, only hypothesized variables of interest, maternal diet, maternal IL-6, and offspring sex, were included in the analysis, as including other factors may have decreased the usable models. Notably, there were no significant differences between the sex (male & female) and post-weaning diet (WSD & CTR) of subjects (post-weaning diet *p*=0.68; sex *p*=0.21) or maternal percent body fat (diet *p*=0.98; sex *p*=0.33). Regardless, these factors and the inclusion of other inflammatory cytokines or glucose/insulin levels would be important to explore in future studies as they may all independently relate to different facets of neurodevelopment.

With the variables considered in this study the use of the Mplus statistical software ensured that only statistically trustworthy models were examined as models that contained issues with data convergence, having a non-positive covariance, negative variance, linear dependencies between variables or a correlation greater than one in the output were filtered out based on error messages produced by the software. This is often missed when using other software, as issues can still exist in a model that ran successfully.

Finally, a considerable portion of the subjects did not have data for all time points as a result of the longitudinal design and the difficult-to attain species. However, this is common in longitudinal designs, and the use of the full information maximum likelihood estimator has been addressed extensively in prior literature ^90–99^.

## Conclusions

Findings from this report improve our understanding of typical macaque cortical development from 4-36 months of age. We were able to characterize the best-fitting functional model for each cortical grayordinate, describing the unique developmental trajectories throughout the brain. Our results emphasize the importance of running these types of analyses on a grayordinate scale, using a longitudinal model. This approach allowed us to observe the uniquely dynamic nature of how the zones surrounding the center of an ROI develop over time. As a proof of concept, we were also able to test how maternal environmental factors diet and interleukin-6 related to the developmental trajectories defined in this study. We found that the described ablation and accumulation zones may be the most susceptible to environmental factors as opposed to the center of a given ROI. The significant clusters associated with the maternal factors displayed not only strong bi-hemispheric tendencies but also showed above-expected concentrations in functional networks previously linked to maternal inflammation in the human literature. The current findings come at a critical time as we continue to understand the developmental costs that accompany the vast increases in obesity, stress rates, and intake of a Western-Style diet in developed countries. Hopefully, we will be able to utilize some of these insights as we learn more about the unknown consequences that may be associated with the increased inflammatory responses associated with the SARS-COV-2 virus that has taken the world by storm.

## Materials and Methods

### 4.1 Macaque Study Overview

This study used Japanese macaques from a well-defined NHP model studying the effects of a Western-style diet (WSD) in relation to a control diet (CTR) ^59, 60, 100^. Maternal and offspring phenotypes have been characterized and described in prior reports ^59, 60, 100–103^. This study and procedures have been approved by the Oregon National Primate Research Center (ONPRC) Institutional Animal Care and Use Committee and also follow the National Institutes of Health guidelines on the ethical use of animals (Figure 1A).

### 4.2 Subjects

Adult females consumed either a CTR diet (Monkey Diet no. 5000; Purina Mills) or a WSD (TAD Primate Diet no. 5LOP, Test Diet, Purina Mills). Diet groups were determined 1.2-8.5 years before offspring birth (age at offspring birth [mean (M) ± SEM]: CTR M= 11.33 ± 3.23 years; WSD M= 8.46 ± 2.31 years). Offspring were housed with their mothers until weaning age (∼8-months) and then socially housed with peers (6-10 other juvenile) and 1-2 unrelated female adults. Further offspring rearing and diet characteristics have been detailed in a recent publication ^59^.

The current study used 53 subjects (Female n=25; CTR n =26), of which the majority (n=44; Female n=19; CTR n =21) of offspring consumed a CTR diet as opposed to a WSD post-weaning. Maternal IL-6 levels were collected from plasma during the 3^rd^ trimester (48.16 ± 9.26 days) prior to offspring birth (maternal age: 9.25 ± 2.95 years). A lower limit of quantification (LLOQ) of 1.23 pg/mL threshold was used for IL-6 levels, excluding subjects below this value. This resulted in a total, 42 (Female n = 19; CTR n = 17) subjects with usable IL-6 measurements. IL-6 concentrations were logarithmically transformed across subjects to center potential outliers and normalize the distribution. Comprehensive plasma collection and analysis procedures have previously been described in detail ^104^.

### 4.3 MRI acquisition

MRI scans were acquired on a Siemens TIM Trio 3 Tesla scanner in a 15-channel knee coil adapted for monkey use. Longitudinal scans occurred at 4 (M = 4.37 ± 0.05), 11 (M = 11.09 ± 0.04), 21 (M = 21.11 ± 0.05) and 36 (M = 36.53 ± 0.09) months of age.

Monkeys were sedated with a single dose of ketamine (10-15mg/kg) for intubation before each scan, followed by a <1.5% isoflurane anesthesia for the rest of the scan. Respiration, heart rate, and peripheral oxygen saturation were monitored throughout.

However, the scan acquisition was adjusted partway through the study to optimize future outputs. For this reason, some scans were acquired using scan acquisition # 1 (n=80), while other scans were acquired with scan acquisition #2 (n=56). While this was a limitation, we believe the outcome was negligible as mean cortical thickness (Supplemental Figure 1) was not significantly different between the two acquisitions while controlling for age (F(1,133)=2.22, *p*=0.139), and thus not included in our models.

A total of four T1-weighted anatomical images were collected for each subject either using acquisition #1 (TE= 3.86 ms, TR= 2500 ms, TI= 1100 ms, flip angle= 12°, 0.5 mm isotropic voxel) or acquisition #2 (TE= 3.33 ms, TR= 2600 ms, TI= 900 ms, flip angle= 8°, 0.5 mm isotropic voxel). Additionally, one T2-weighted anatomical image was acquired for acquisition #1 (TE= 95 ms, TR= 10240 ms, flip angle= 150°, 0.5 mm isotropic voxel) or acquisition #2 (TE= 407 ms, TR= 3200 ms, 0.5 mm isotropic voxel). Additional scan types were acquired in these sessions, but not used for this manuscript.

### 4.4 MRI preprocessing

The current study used a macaque-specific modified version of the human connectome project (HCP) minimal preprocessing pipeline ^105^. Details on preprocessing have previously been described in our prior publication ^58^. In brief, this pipeline was built to analyze outputs in surface space and used FMRIB Software Library (FSL) ^106–108^, the FreeSurfer image analysis suite (http://surfer.nmr.mgh.harvard.edu109,110, Advanced Normalization Tools (ANTs) (version 1.9; http://stnava.github.io/ANTs/) in combination with in-house established tools. The general structure follows the HCP stages ^105^, with macaque-specific modifications occurring before and in the PreFreeSurfer, FreeSurfer, and PostFreeSurfer stages. Subjects were aligned to the Yerkes19 atlas space ^111^. Finally, this modified pipeline follows previously established standards for human data and the ABCD project (available here https://github.com/DCAN-Labs/ or here https://github.com/ABCD-STUDY/abcd-hcp-pipeline) and has been prepared for a similar NHP release (https://zenodo.org/record/3888969).

### 4.5 General analysis overview

Cortical thickness development was investigated in relation to maternal diet, IL-6 during gestation, and offspring sex using latent growth curve (LGC) analyses. Growth trajectories over time and related predictors/covariates were estimated with LGCs, which stem from the structural equation modeling (SEM) framework ^68–70^. An in-depth description of how LGC models work for this type of analysis has been detailed in our previous paper (“Maternal interleukin-6 is associated with macaque offspring amygdala development and behavior”:^58^). In brief, first, the best-fitting model to characterize the typical growth trajectory is established as the unconditional model. Then predictors and covariates are added to the model to see how they relate to the different aspects of the typical growth trajectory (e.g., the starting point [intercept] or the growth [slope]). The best-fitting model is determined by testing out the simplest model (intercepts only), followed by more complex models (intercept & slope, or intercept, slope & quadratic). A chi-squared difference test is used to see if adding the additional functional forms significantly improves the model. The simplest model that isn’t significantly improved by additional functional form/s is considered the best-fitting model ^112^ (Figure 1B).

Individual models were analyzed using version 8 of Mplus ^113^, with the robust maximum likelihood estimator for non-normal data and full information maximum likelihood for missing data ^95^. A large body of research supports the use of this method in studies with missing data at different time points ^90–97^. Best-fitting models were identified using in-house derived code in MATLAB (The MathWorks, Inc., Natick, Massachusetts, United States), and the procedures of what this code did are explained throughout the remainder of the methods and depicted in Figure 1B. Finally, figures were created using Jupyter Notebooks, MATLAB, PowerPoint, and the Connectome Workbench visualization software ^114^.

### 4.6 Characterizing typical cortical thickness development on a grayordinate level

To establish typical cortical thickness development across the four time-points, the previous convention in the field ^67^ was followed, which determined the best-fitting unconditional model on every surface grayordinate (n=56522) (Figure 1B).

Consequently, for each grayordinate, multiple latent growth models using different functional forms (intercept, slope, quadratic, and splines on the 4, 11, 21, and 36-month time points) were computed based on prior convention ^67^. Chi-squared difference tests were then computed between the most simple (intercepts only) and more complex models (e.g., intercept & slope) to determine the best-fitting model of each grayordinate (Figure 1B). This enabled characterization of the best-fitting growth trajectory of each grayordinate and also allowed us to observe how the estimated mean cortical thickness of each grayordinate changed across time. For visualization purposes, a movie is provided (Supp Mov. 1) of the estimated mean cortical thickness development from 4 to 36-months of age. The thickness at time points (in months) between actual scan dates was interpolated using a linear interpolation for change of thickness in each month.

### 4.7 Introducing predictors to the unconditional models

After modeling typical cortical thickness development, predictors of interest (maternal IL-6, maternal diet, offspring sex) were introduced. This allowed us to identify how each predictor related to the latent growth variables (i.e., intercept, slope, quadratic) of each grayordinate trajectory while controlling for the impact of the other predictors. Estimates of the relationships between predictors and growth variables can be displayed on the brain for visualization purposes (Figure 4-Figure 6).

### 4.8 Cluster detection analysis

Having established estimates, p-values, and standard errors for each model, we next identified which specific regions from this analysis were most notable using a cluster detection analysis. To avoid issues with multiple comparisons and identify the most important areas, these estimates were standardized and run through a cluster detection algorithm based on previously defined guidelines ^72^. Clusters of voxels defined by estimates and standard errors were ordered by size, permutated, and top clusters were kept based on a predefined z-score threshold. In this case, clusters above/below a z-score of 2.36 were used, yielding a corrected significance p-value of 0.01 in line with previous findings using cortical thickness data ^115^.

### 4.9 Cluster surface area within functional networks

To increase generalizability, the surface area of each cluster was calculated and grouped by functional networks. Networks and ROIs used for this have been previously described ^74, 77^. A chi-square test was used to calculate if the observed clusters significantly differed from the expected [*(expected-observed)^2^/expected*] for each network (Figure 7). The expected was calculated by obtaining the ratio of observed total cluster surface area for each predictor to the total cortical surface area (7649.77 mm^2^) and multiplied by the surface area of each network (Default [DMN]: 2026.28 mm^2^, Visual [VIS]: 2211.05 mm^2^, Limbic [LIM]: 793.65 mm^2^, Dorsal Attention [DAN]: 748.34 mm^2^, Insular-Opercular [INO]: 322.47 mm^2^, and Auditory [AUD]: 150.77 mm^2^) (Figure 7)

### 4.10 Testing how maternal diet, IL-6, and offspring sex relate to whole-brain average cortical thickness

Finally, to test whether findings were not just a function of changes in overall total cortical thickness, this same analysis was repeated on mean total brain cortical thickness development as opposed to individual vertices across the cortex. This was run as proof of concept that the more detailed grayordinate specific approach added information and was thus needed to identify how predictors related to changes in specific aspects of cortical development depending on the region as opposed to the total brain. To do this, the best-fitting unconditional model of typical total brain development was identified using a chi-squared difference test. Then, predictors of interest (IL-6, Maternal Diet, Sex) were added to see how they related to the functional forms (intercept and slope) of this new conditional model (Supplemental Figure 4, Supplemental Table 1).

## Data Availability

The data supporting these findings are available upon reasonable request from the corresponding author

## Code Availability

The code used for the analysis of this study is available upon reasonable request to the corresponding author. Data were processed with our publicly available preprocessing pipeline accessible at Zenodo (https://zenodo.org/record/3888969) or docker ( https://hub.docker.com/r/dcanlabs/nhp-abcd-bids-pipeline)

## Acknowledgments

This work was supported by National Institutes of Health (grants R01MH107508 to E.L.S.; R01 MH096773 to D.A.F.; R01 MH115357 and U01 AA021691 to D.A.F), the Masonic Institute for the Developing Brain (to D.A.F.), the Lynne and Andrew Redleaf Foundation (to D.A.F.)

## Author Contributions

Conceptualization, D.A.F, E.L.S., J.S.B.R; project supervision, D.A.F, E.L.S., T.X, E.F, A.M.G., M.P.M; data acquisition and processing, J.R.T., J.L.B., D.S., J.S.B.R., A.M., J.Y.Z., S.P., M.B., Analyses and methodology, J.S.B.R., R.H., A.G., E.T., E.S., O.M.D., E.F., writing-original Draft, J.S.B.R.; writing-review & editing, J.S.B.R. D.A.F, T.X, E.L.S., R.H., A.M.G., A.M., E.T., O.M.D., M.P.M.

## Materials & Correspondence

Correspondence to Damien A. Fair

