## Supplemental Materials for "Vertex-wise characterization of Non-Human Primate cortical development with prenatal insights"

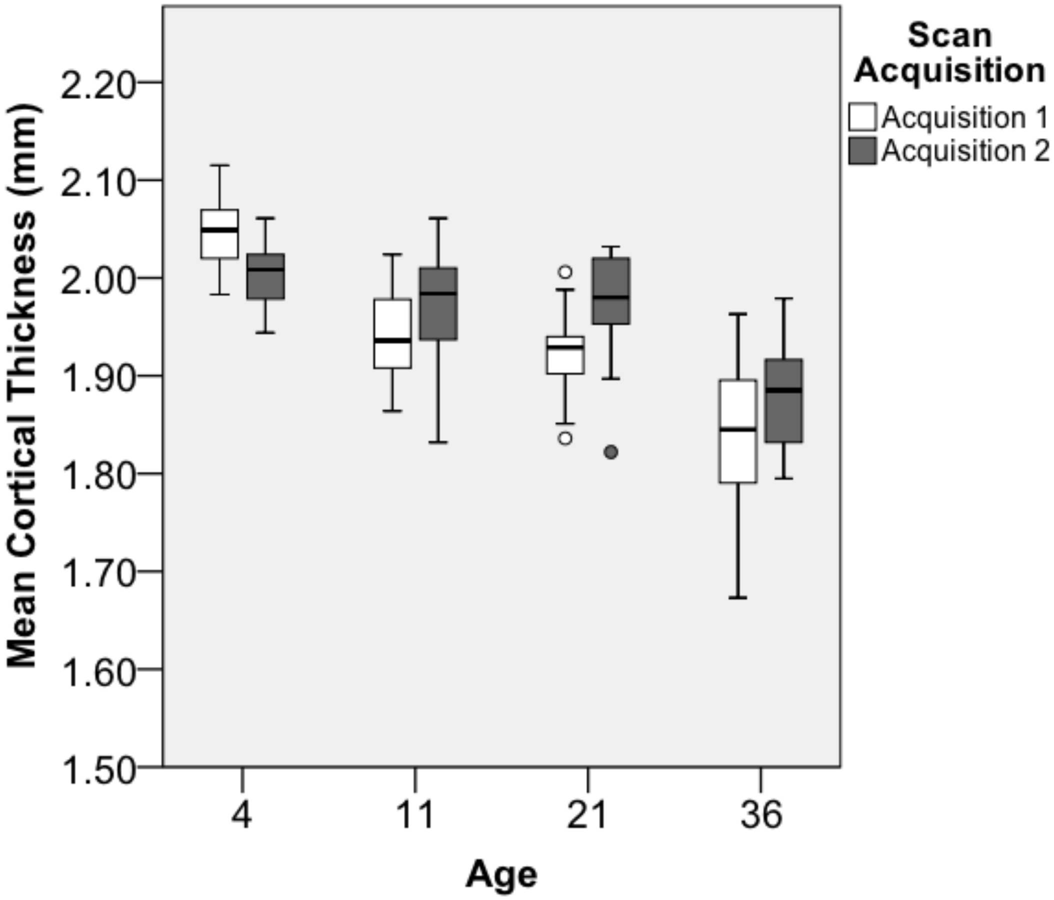


Supplemental Figure 1: Comparing the scan acquisition across age to show they were not significantly different.

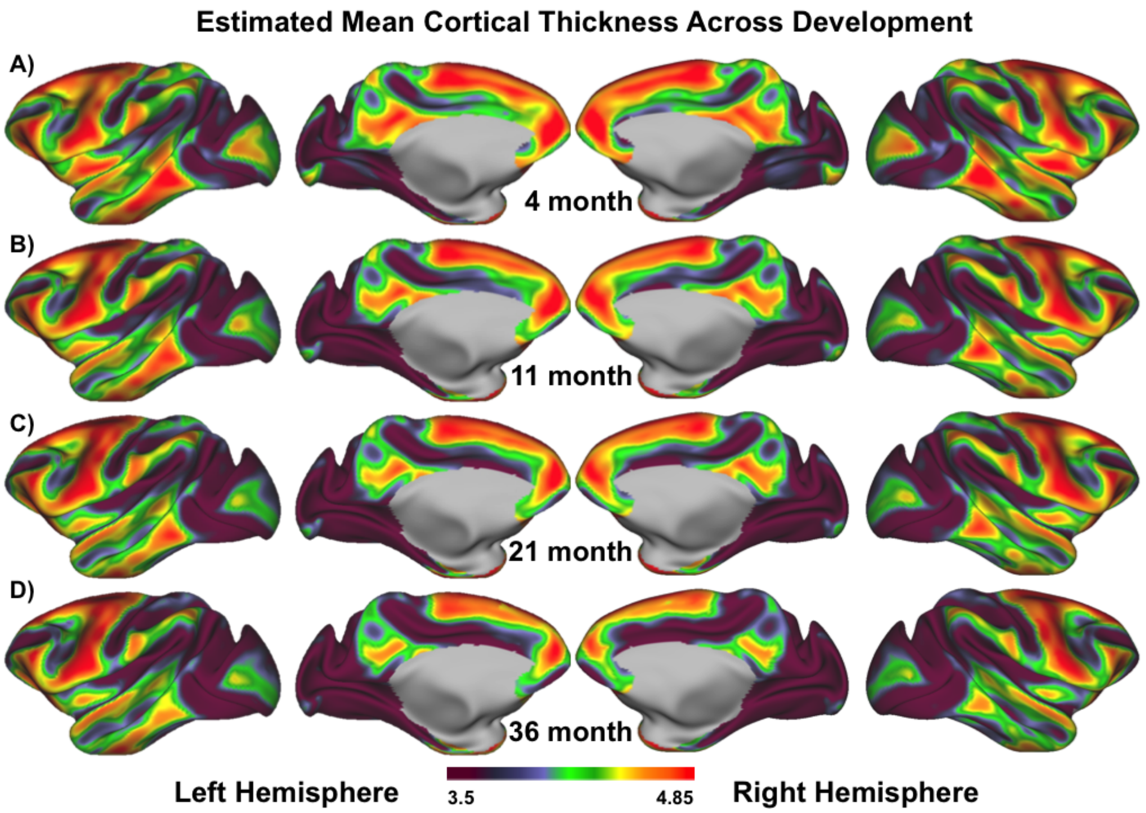


Supplemental Figure 2: Both hemispheres of mean cortical thickness

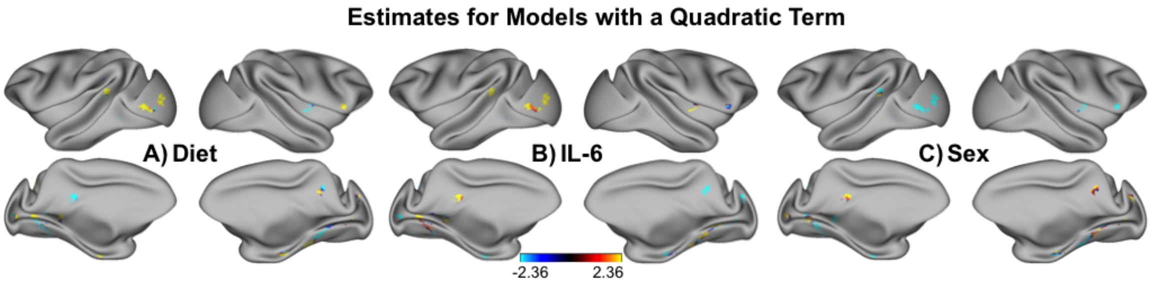


Supplemental Figure 3: Shows the estimates for the quadratic term for maternal diet (A), maternal IL-6 (B), and offspring sex (C). As the majority of grayordinates were not described by a best-fitting quadratic trajectory (Figure 1.3), only a few regions contain this quadratic estimate.

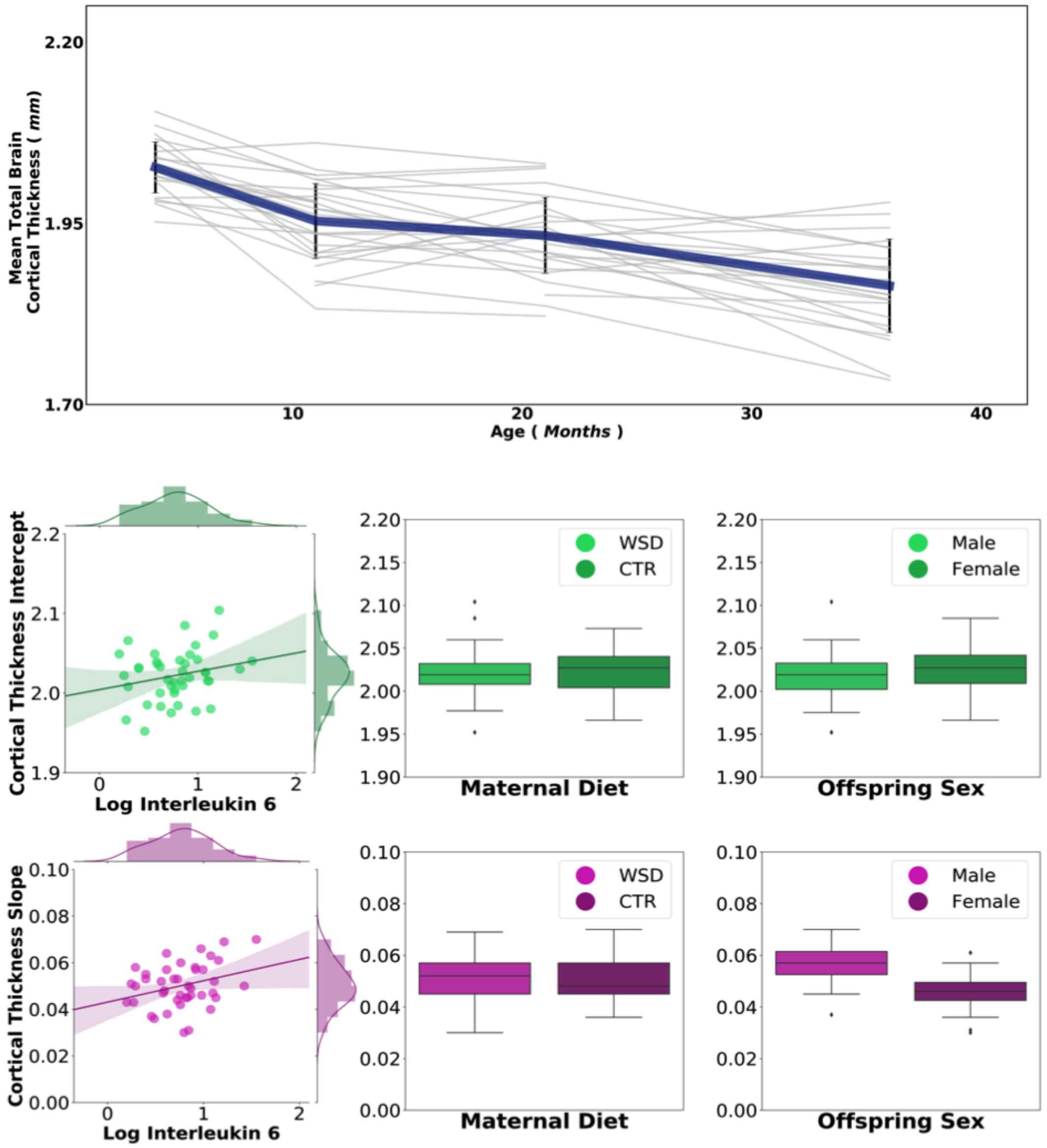


Supplemental Table 1: Model statistics for average total brain cortical thickness

| | Unconditional Model | | Conditional Model | |
|---|---|---|---|---|
| Parameter | Estimate | S.E. | Estimate | S.E. |
| Intercept Mean | *** 2.019 | 0.007 | ***2.001 | 0.020 |
| Intercept Variance | ***0.002 | <0.001 | **0.001 | <0.001 |
| Slope Mean | ***0.051 | 0.005 | ***0.047 | 0.012 |
| Slope Variance | <0.001 | <0.001 | <0.001 | <0.001 |
| Intercept & Slope Covariance | <0.001 | <0.001 | <0.001 | <0.001 |
| Predictors of Intercept | | | | |
| Interleukin-6 | N/A | N/A | 0.024 | 0.022 |
| Maternal Diet | N/A | N/A | -0.005 | 0.017 |
| Offspring Sex | N/A | N/A | 0.010 | 0.016 |
| Predictors of Slope | | | | |
| Interleukin-6 | N/A | N/A | 0.008 | 0.012 |
| Maternal Diet | N/A | N/A | 0.008 | 0.009 |
| Offspring Sex | N/A | N/A | ł -0.015 | 0.009 |

Note: ł= *p* <.10; **p*<.05; ***p*<.01;****p*<.001. N/A *indicates covariates not included in the final model*

Sex Findings:

*How offspring sex, related to cortical thickness development.*

Offspring sex was included in the model as a covariate, allowing us also to identify how offspring sex was associated with cortical thickness development. Associations with offspring sex were seen relating to the 4-month starting point (intercept) and the rate of cortical thickness growth (slope) across the four time-points

(Figure 6). Due to the small sample size and number of comparisons run, no sex*diet or sex*IL-6 interactions were added to the model, so sex was only treated as an independent predictor in the model alongside IL-6 and Diet. Males were associated with increased bilateral starting (intercept) cortical thickness in the dorsal part of the visual anterior cortex, as well as the inferior parietal cortex (Figure 6A&C; [green clusters; yellow arrows]). However, males also showed decreased starting cortical thickness in the orbitomedial prefrontal cortex (Figure 6A&C; [green clusters; blue arrows]).

Regarding the rate of change, males exhibited increases in the superior parietal cortex and the primary somatosensory cortex (Figure 6B&C; [pink clusters; yellow arrows]). Finally, there was a unique relationship between sex and the shape of development (quadratic). Males exhibited bilateral decreased quadratic clusters in the hippocampus (Figure 6C& Supplemental Figure 3; [white clusters; blue arrows]). These clusters also followed the previously described trend of falling on the edge of regions that had either the thickest or thinnest cortical thickness, as can be observed in the superior parietal/primary somatosensory cortex clusters and the orbitomedial prefrontal cortex clusters, for example.

Finally, sex network findings were distributed throughout the brain. However, sex findings only significantly diverged from the expected in the growth characteristics of cortical thickness development (i.e. slope and quadratic) but not the intercept (4-month) results. Sex was associated with higher than expected surface area in the VIS (Slope: $\chi^2$ (1) = 24.63, p > .001; Quadratic: $\chi^2$ (1) = 8.40, p > .01) and AUD (Slope: $\chi^2$ (1) = 38.31, p > .001; Quadratic: $\chi^2$ (1) = 7.61, p > .01) networks for the slope and quadratic results, with lower than expected surface area in the DMN ($\chi^2$ (1) = 6.40, p > .05), LIM ($\chi^2$ (1) =

7.78, p > .01), and INO ($\chi^2$ (1) = 6.04, p > .05) networks for the slope results. Additionally, there were lower than expected quadratic surface area results in the DAN ($\chi^2$ (1) = 7.30, p > .01) and SMN ($\chi^2$ (1) = 9.03, p > .01) networks (Figure 7).

*Findings were region-specific rather than a global phenomenon.*

Finally, to show that these types of analyses are region-specific and do not just reflect global changes across the brain, we modeled mean whole-brain cortical thickness development in relation to maternal diet and IL-6. Similar to the individual grayordinate models, the development of average total brain cortical thickness was assessed first to determine the unconditional model before introducing predictors to establish the conditional model (Supplemental Figure 4, Supplemental. Table 1). A spline model freeing the 11-month time point best described the unconditional mean cortical thickness development ($\chi$2 (5) = 9.20, *p* = 0.101, CFI = 0.81, TLI = 0.77, RMSEA = 0.13). Introducing the predictors and covariates (Maternal Diet, Maternal IL-6 and Offspring Sex) to the model improved the model fit ($\chi$2 (11) = 12.64, *p* = 0.317, CFI = 0.86, TLI = 0.77, RMSEA = 0.06) but failed to show any significant relationships between variables IL-6 (Intercept: *B*=0.024, *p*=0.281, Slope: *B*=0.008, *p*=0.515), diet (Intercept: *B*=-0.005, *p*=0.775, Slope: *B*=0.008, *p*=0.376), or sex (Intercept: *B*=0.010, *p*=0.513, Slope: *B*=-0.015, *p*=0.095). These findings are depicted in Supplemental Figure 4 and Supplemental Table 1.

As hypothesized, using mean cortical thickness was too crude of a measure and did not capture the complicated nature of how maternal IL-6 and diet relate to cortical thickness development. A grayordinate level analysis would allow for region-specific bidirectional effects as opposed to the outcomes from a whole-brain approach.

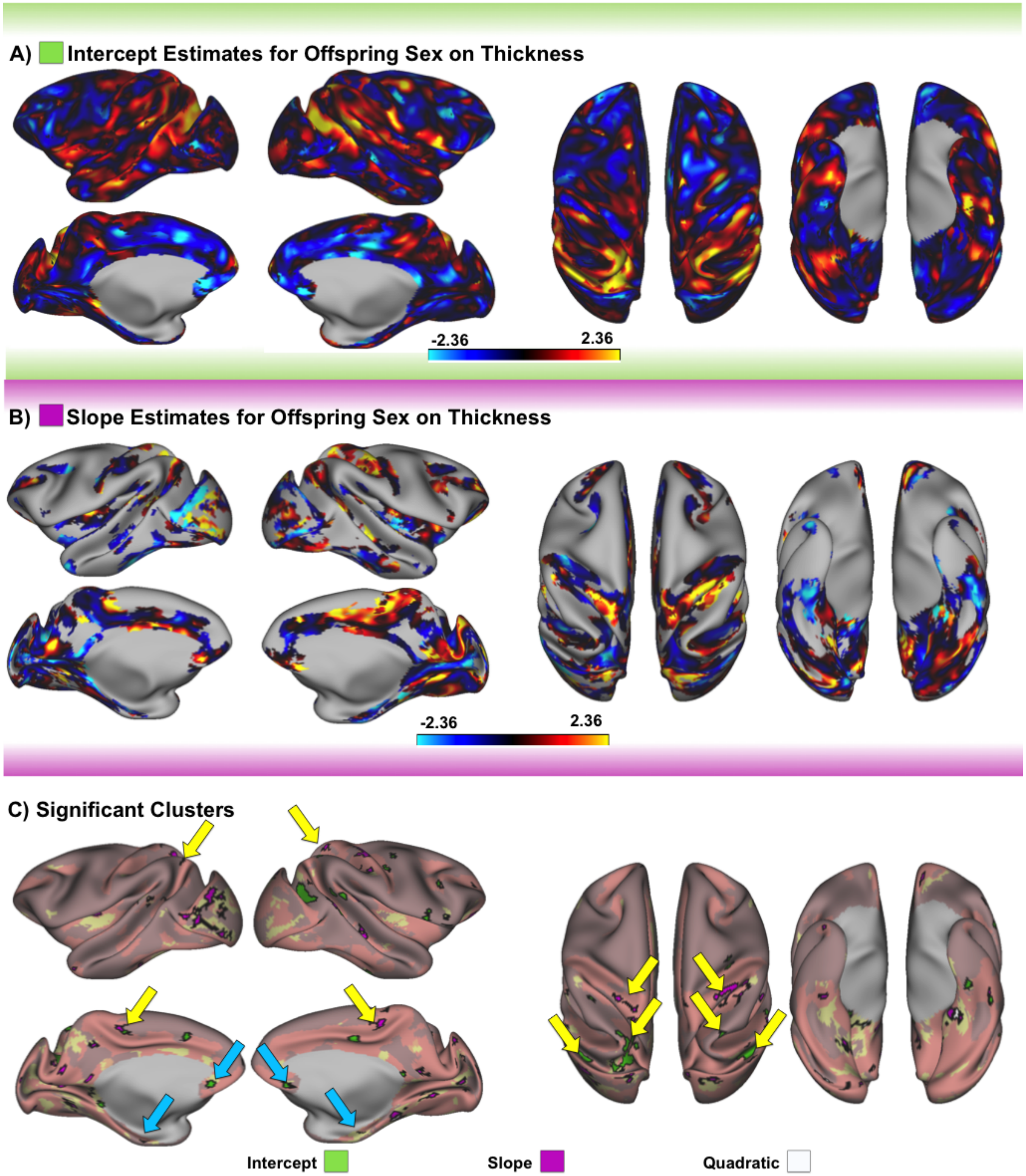

A) Intercept Estimates for Offspring Sex on Thickness
-2.36
2.36
B) Slope Estimates for Offspring Sex on Thickness
-2.36
2.36
C) Significant Clusters
Intercept
Slope
Quadratic

Figure 6: The influence of offspring sex on cortical thickness development. (A) Indicates the estimates of how sex relates to the intercepts of the models (B) shows the estimates of how sex relates to the slope parameter, and (C) highlights the significant clusters from this analysis. Green clusters are clusters where sex had a significant relationship with the intercept, and pink clusters show the significant slope clusters. White clusters indicate the quadratic; however, since there were so few best-fitting quadratic models, the estimates are reported in the supplemental materials. Arrows highlight some interesting bilateral clusters where males were associated with decreased (blue arrow) or increased (yellow arrow) intercept (green cluster) or slope (pink cluster). The direction of association (highlighted by the color of arrows) can be identified by the color of the estimates (A&B).

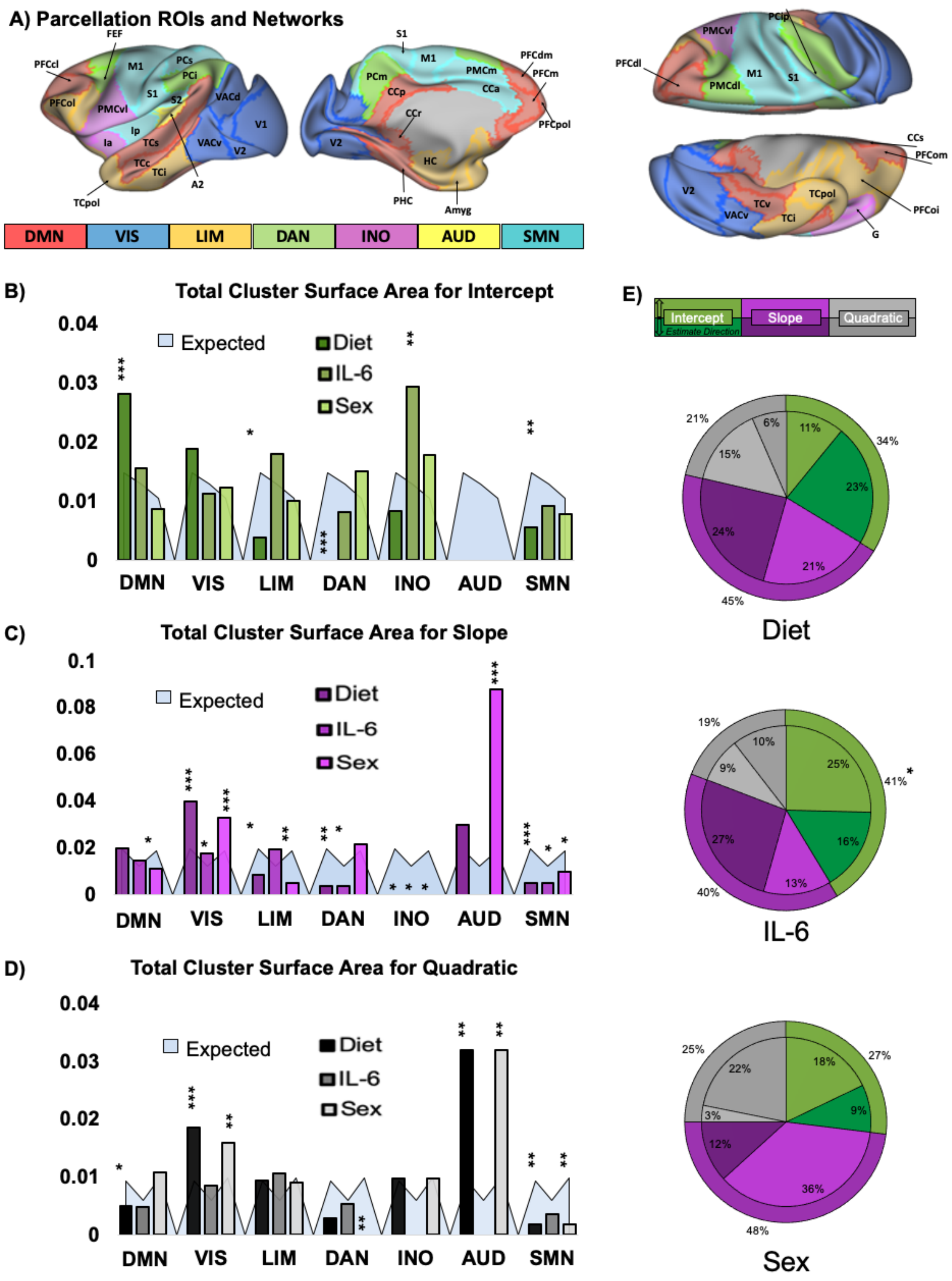

A) Parcellation ROIs and Networks
DMN
VIS
LIM
DAN
INO
AUD
SMN
B) Total Cluster Surface Area for Intercept
Expected
Diet
IL-6
Sex
C) Total Cluster Surface Area for Slope
D) Total Cluster Surface Area for Quadratic
E)
Intercept
Slope
Quadratic
Estimate Direction
Diet
IL-6
Sex

Figure 7: Summary results describing the cluster surface area trends. (A) Shows the functional networks and ROIs (Bezgin et al. 2012; Grayson et al. 2016). (B-D) highlight the surface area of the significant clusters for the intercept (B) slope (C) and quadratic (D). These graphs show the significant clusters for each predictor (Diet, IL-6, and Sex) and where, in relation to the parcellation, these clusters fell, as indicated by the network they are in. The total cluster surface is plotted on top of the expected surface area for each network and predictor (light blue). Finally, the pie charts (E) indicate the percent of the surface area of the clusters related to the intercept (green), slope (pink), or quadratic (grey), with lighter colors indicating a positive (i.e., the predictor is associated with increased thickness) and darker a negative (i.e., the predictor is associated with decreased thickness) association. Note: $*P<0.05$; $**P<0.01$; $***P<0.001$. .

*Sex-specific findings also relate to sex-specific results in the literature.*

Sex differences in cortical thickness development have been heavily studied (Raznahan et al. 2011; Lyall et al. 2015; Vijayakumar et al. 2016; Gennatas et al. 2017). Sex differences observed in this study match up to current literature in humans (Amlien et al. 2016; Gennatas et al. 2017). The current studies macaque timeline of 4 to 36-months of age roughly translates to 1 year through early puberty in humans, generally speaking. However, individual processes may be expedited or delayed depending on the developmental species-dependent function and anatomy. For example, interspecies cortical expansion is lower in the occipital and temporal regions between macaque and human development, with the interspecies expansion maps correlating more strongly during early (4-10yrs) than late (17-30yrs) development (Amlien et al. 2016). The current data showed that males predominantly had thicker occipital cortical thickness starting points (intercept) but then a general increased rate of cortical thinning over time (slope). This occipital trend heuristically matches up with developmental findings showing increased male thickness in early ages (up until around 15 years), with decreased cortical thickness in males later in life (up until early adulthood) (Gennatas et al. 2017). Parietal cortices were also, in general, thicker in males than females, with decreased frontal thickness in males compared to females (Gennatas et al. 2017). It is important to note that the macaque and human studies do not match in terms of homologous developmental timelines. This is a limitation as a result of no studies existing in the human literature that investigate sex differences in cortical thickness from 1 year to early puberty. Despite this limitation, it is reassuring to see that the general trends observed in the human literature match up to the most part with the current findings.
